# A Multi-Modal Deep Learning Framework with Both Sequence and Structure for Tumor Antigens Prediction

**DOI:** 10.1101/2024.11.06.622193

**Authors:** Ruofan Jin, Jingxuan Ge, Guanqiao Zhang, Ziyan Deng, Kim Hsieh, Tingjun Hou, Ruhong Zhou

## Abstract

Tumor antigens are key targets in cancer immunotherapies that can be recognized by T cell receptor and induce immune responses. However, precision screening of immunogenic tumor antigens remains a great challenge due to human leukocyte antigen (HLA) restriction and tumor antigen escape. Here, we introduce MultiTAP (Multi-modal Tumor Antigen Predictor), a pioneering multi-modal framework with TCR-peptide-HLA sequence and structure features incorporating an attention mechanism designed to accurately identify tumor antigens with immunogenic properties. By constructing the multi-modal TCR-peptide-HLA Dataset (TPHD) and integrating its sequence and structure, we perform antigen feature enhancement using peptide-HLA (pHLA) structural features at the residue level, achieving interpretable prediction of immunogenicity for tumor antigens. Relative to existing baseline models, the MultiTAP framework has exhibited superior efficacy in predicting the immunogenicity of tumor antigens. Through comprehensive out-of-distribution (OOD) assessments, MultiTAP has maintained predictive robustness across diverse HLA phenotypes and the continuously evolving landscape of epitope distributions. Overall, MultiTAP presents a brand-new and promising approach for cancer immunotherapies that target tumor antigens.

## Main

Endogenous tumor antigens have emerged as pivotal targets in cancer immunotherapy^1,2,3^, garnering significant interest within the scientific community^4,5^ and becoming a focal point of extensive research efforts^6,7,8^. These antigens are presented on the cell surface by MHC class I molecules in the tumor microenvironment and subsequently recognized by CD8+ T cells, initiating tumor-specific immune responses and facilitating T cell-mediated tumor cell elimination^9,10,11^. Originating primarily from tumor-specific somatic mutations and absent in normal tissues, these antigens exhibit unique immunogenicity that effectively activates the immune system and elicits robust T cell responses^12^. Immunotherapeutic strategies targeting these neoantigens have shown promise in augmenting the antitumor efficacy of existing treatments, particularly in the development and implementation of personalized therapeutic approaches^13^. Consequently, precise identification and screening of endogenous tumor antigens are fundamental and critical for the advancement of cancer immunotherapy.

Among the vast array of mutated peptides, only a limited subset can elicit potent antitumor immune responses^14^. To enable efficient screening of endogenous tumor antigens, researchers have employed computational methods using immunogenicity datasets of TCR-recognizable antigens to build predictive models^15^. The CDR3β region of the TCR, known for its highest diversity and primary role in antigen recognition, is typically considered the most critical contributor to peptide binding^16,17,18^, leading many studies to focus solely on TCR CDR3β sequence information to examine the specificity of TCR-peptide interactions. For instance, ImReX^19^ employs a novel feature representation approach based on the pairwise combination of physicochemical properties of the amino acids in the CDR3 and epitope sequences, combined with a convolutional neural network (CNN) architecture for TCR-epitope recognition evaluation. Similarly, ERGO-I^20^ introduces a sequence representation method grounded in natural language processing (NLP) to predict binding between TCRs and peptides. In addition, PanPep^21^ has demonstrated high efficiency in TCR recognition across different peptide types by integrating meta-learning and the neural Turing machine framework. Furthermore, TEINet^22^ utilizes transfer learning to predict TCR binding specificity. Recent advancements such as ERGO-II^23^ and NetTCR-2.0^24^ have underscored the critical role of TCR CDR3α in antigen recognition, demonstrating that the inclusion of CDR3α not only enhances the understanding of TCR-antigen interactions but also expands the molecular relationship information in this process. This shift reflects a growing trend to integrate structural biology insights and biophysical data into TCR-antigen interaction predictions, ultimately improving the computational methods for assessing the immunogenicity of antigens.

While efforts have been made for accurate screening of TCR-recognized antigens, it is evident that current models encounter some challenges in effectively and reliably predicting the immunogenicity of rare epitopes^25^. It is primarily because most rare epitopes are endogenous tumor antigens^26,27^, which become recognizable by TCRs only after being presented on the tumor cell surface by HLA, subsequently inducing T cell-mediated immune responses that promote tumor cell apoptosis^28^. However, HLA phenotype limitations and reduced HLA expression in tumors further hinder the effective presentation of these endogenous tumor antigens^29,30,31^. Such challenges have inspired researchers to develop computational methods that bridge from antigen presentation to TCR–antigen recognition, incorporating HLA-related features to enhance their generalization capabilities for rare or novel epitopes^25^. Yet, current approaches primarily consider HLA pseudo-sequence^32,33^, which neglects crucial amino acid information within full-length sequences. Consequently, these methods fail to account for functional variations and specificities of HLA alleles, leading to a lack of comprehensive understanding of the complexity and diversity of HLA molecules, as well as insufficient information on how different HLA phenotypes influence epitope presentation^34^. Such limitations ultimately affect the accuracy of immunogenicity predictions for endogenous tumor antigens.

Existing studies have established that the crystal structure of HLA molecules is critical for both tumor antigens presentation and their immunogenicity^10,35,36^. Notably, different HLA alleles exhibit significant structural differences in their binding affinities for specific peptides^37,38^, which simultaneously affects the specificity of TCR interactions with these antigens^39^. However, current computational models have yet to effectively incorporate HLA structural information for the accurate prediction of the immunogenicity of tumor specific antigen^32,33^. Recent investigations, such as those conducted by ITN^40^ and TransfIGN^41^, have revealed that the inclusion of HLA structure and its associated characteristics in computational frameworks markedly improves the precision of antigen presentation predictions. Consequently, the exploration of HLA molecular structure and its relevant features within computational TCR-antigen recognition models merits further research and scrutiny.

Here, we introduce MultiTAP (Multi-modal Tumor Antigen Predictor), an innovative multi-modal model incorporating an attention mechanism^42^. MultiTAP integrates TCR-peptide-HLA sequences and peptide-HLA (pHLA) structural data to construct a predictive model for identifying endogenous tumor antigens. We have developed a multi-modal dataset, namely the TCR-peptide-HLA Dataset (TPHD), which encompasses three major databases: IEDB^43^, McPAS-TCR^44^, and VDJdb^45^, and obtained thousands of binary complex structural data of pHLA utilizing the xTrimo-Multimer model for protein complex structure prediction^46,47^. Supported by TPHD, MultiTAP employs pre-trained model-based methods combined with sequence alignment to extract sequence-derived features^48^. It then merges sequence and structural information through an attention module, forming multimodal features that characterize TCR recognition of antigens presented by HLA, thereby enabling predictions of antigen immunogenicity.

A comparison of MultiTAP with baseline models on the same dataset demonstrated superior performance in predicting immunogenicity. Furthermore, we proposed clustering-based methods for segmenting antigens and HLA alleles datasets to address two key issues: 1) the model’s ability to recognize antigens across different cellular backgrounds, and 2) its capacity to accurately identify antigens under varying HLA epitope restrictions. Comparative analysis with existing baseline models highlighted MultiTAP’s superior performance in predicting neoantigens and its improved accuracy in assessing antigen immunogenicity under HLA restriction. Exploring the role of multimodal information, particularly pHLA structural data, in the antigens screening process by MultiTAP revealed at the residue level that the integration of HLA presentation information is crucial for precise epitope immunogenicity prediction. MultiTAP provides novel insights and methodologies for investigating tumor antigen immunogenicity, introducing an innovative multi-modal data feature analysis framework that integrates pHLA structures. This approach establishes a crucial foundation for T-cell immunotherapies targeting endogenous tumor antigens.

## Results

### 1. MultiTAP overview

We introduce MultiTAP, a multimodal model that leverages hierarchical features from the TCR-peptide-HLA structure (Figure 1a). The data sources for these features include CDR3-paired TCR protein sequences, antigen sequences, HLA protein sequences, and pHLA binary complex structures (Figure 1b). Due to the limited availability of crystal structure data for pHLA binary complexes, we utilized the xTrimo-Multimer model developed by BioMap to obtain the pHLA structural data.

**Fig. 1:**
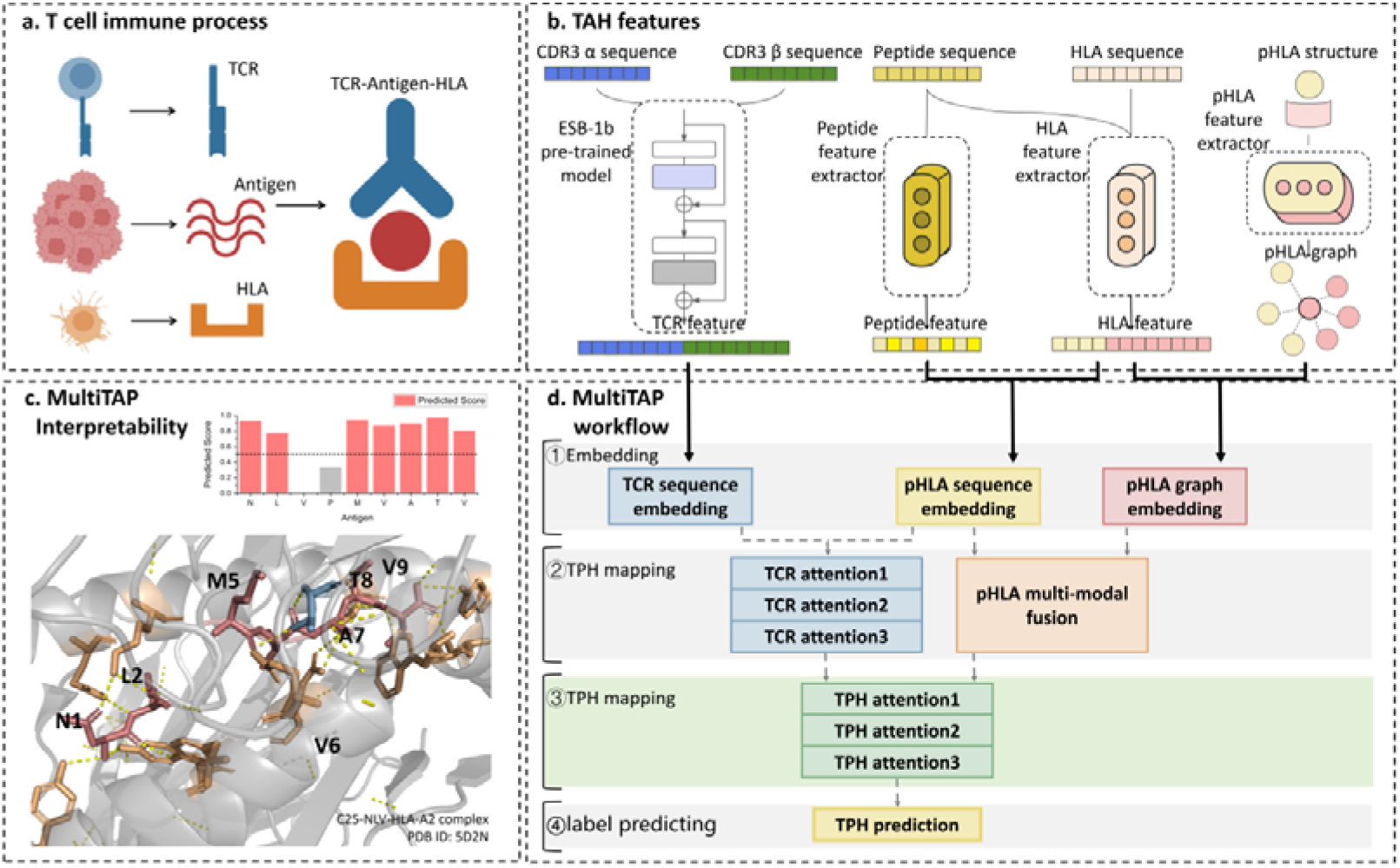
Schematic overview of the MultiTAP model framework and its functional components. **a**, Structural depiction of the TCR-peptide-HLA ternary complex, illustrating the essential interactions underpinning T-cell immunity. **b**, Representation of the input processing pipeline, highlighting the sequence and structural feature generation modules for the TCR-peptide-HLA complex. **c**, Visualization of the attention mechanism within MultiTAP, demonstrating the contribution of each residue to immunogenicity prediction. The analysis reveals at residue level that interactions between the antigen and HLA, as well as between the antigen and TCR, play pivotal roles in antigen immunogenicity prediction. **d**, Workflow of the MultiTAP model, encompassing four primary stages: embedding, peptide-HLA mapping, TCR-peptide-HLA mapping, and label prediction. The mapping stages incorporate attention algorithms for peptide-HLA and TCR-peptide-HLA interaction embedding mapping, enabling residue-level attention scoring and enhancing interpretability.

Leveraging protein pre-trained models, sequence alignment, and structural visualization (Figure 1b), we extracted features across multiple hierarchical levels, encompassing primary to tertiary protein structures, to serve as inputs for the MultiTAP multimodal model. As depicted in Figure 1d, MultiTAP integrates an attention mechanism with a Graph Neural Network (GNN) to amalgamate features derived from both pHLA structural and sequence data, effectively simulating the post-epitope presentation binding state. The model subsequently maps TCR sequence features and pHLA binding state features onto potential epitope sequence features. Through iterative model refinement, it captures the “virtual binding” state of the epitope, which is ultimately utilized to assess TCR binding affinity. This approach enables the prediction of tumor antigen immunogenicity (Figure 1c).

### 2. Multi-modal Data Integration Elevates MultiTAP’s Immunogenicity Prediction

We posit that the multimodal architecture of MultiTAP is pivotal for accurately predicting the immunogenicity of tumor antigen peptides. To substantiate this, we curated a comprehensive multimodal dataset and performed a series of feature ablation studies. It enabled a systematic assessment of each feature’s contribution to the model’s performance, thereby highlighting the significance of integrating diverse data modalities—such as CDR3-paired TCR sequences, peptide sequences, HLA sequences, and structural information of pHLA complexes—in enhancing the precision of antigen immunogenicity predictions.

We aggregated TCR-peptide-HLA binding data from three databases: IEDB^43^, McPAS-TCR^44^, and VDJdb^45^. Then utilized xTrimo-Multimer to generate structural data for pHLA complexes based on pHLA sequences obtained from TCR-peptide-HLA binding data. Employing a random combination approach, we produced fivefold negative sample data, which evaluates the validity of this negative sample generation method. This process resulted in a multimodal dataset, the TCR-peptide-HLA Dataset (TPHD), comprising nearly 17,000 data points (Figure 2a).

**Fig. 2:**
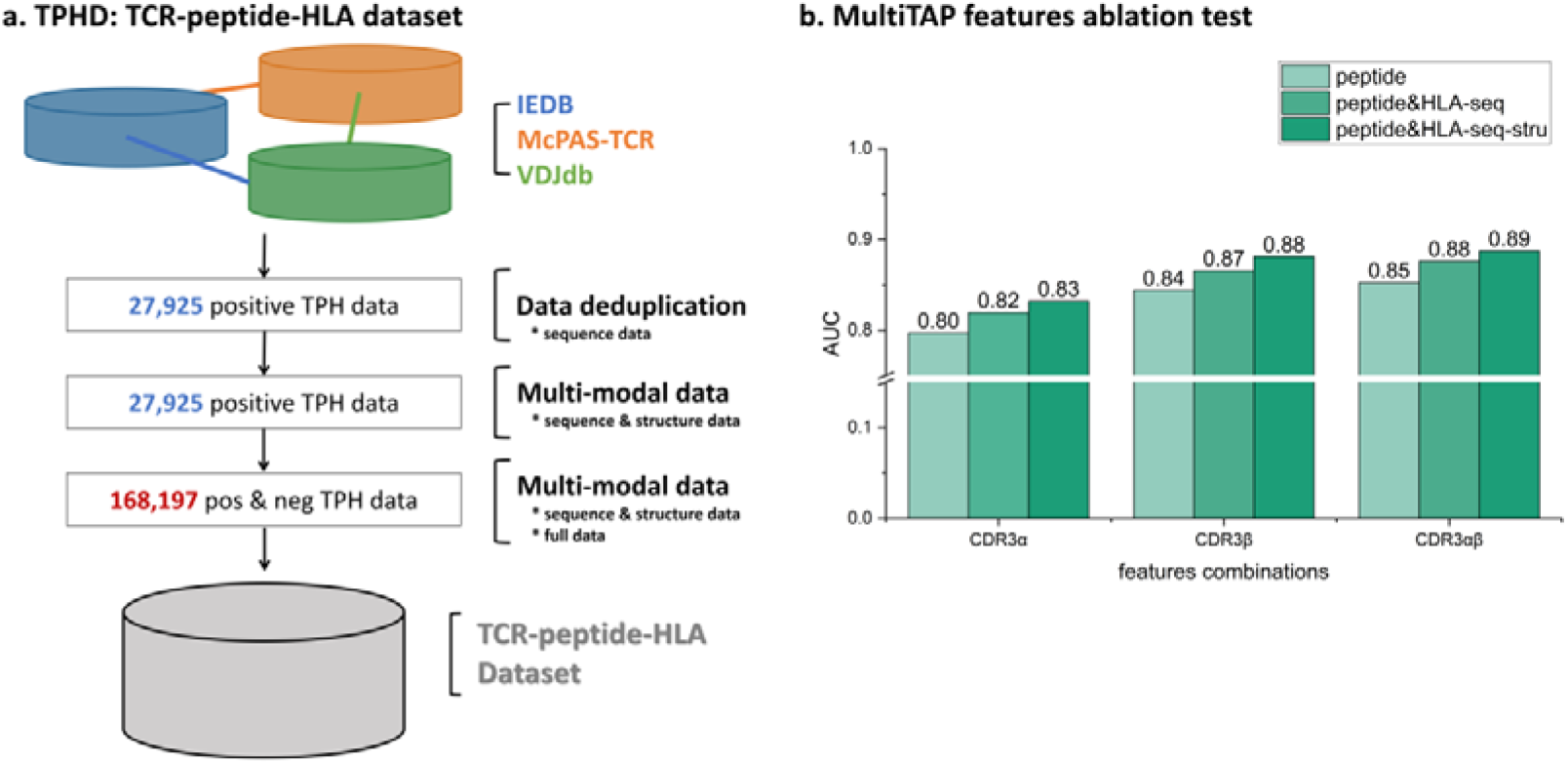
TCR-peptide-HLA Dataset construction and MultiTAP feature ablation experiment. **a**, Overview of the data collection and curation pipeline, integrating data from IEDB, VDJdb, and McPAS-TCR. The selection criteria focused on CDR3-paired data”, “human tumor data” and records containing HLA allele information, resulting in a deduplicated dataset of 27,925 positive TCR-peptide-HLA samples. pHLA binary complex structural modeling was performed using xTrimo-Multimer, producing a multimodal dataset. Through a five-fold randomized cross-sampling method, a comprehensive dataset comprising 168,197 positive and negative samples was generated, completing the construction of the TCR-peptide-HLA dataset (TPHD). **b**, Results from feature ablation experiments assessing the contribution of CDR3α sequences, HLA sequences, and HLA structural data to the predictive performance of MultiTAP, demonstrating their positive impact on prediction accuracy.

To evaluate the impact of various feature combinations on model performance, we derived six distinct feature sets from the TPHD dataset: 1) TCR CDR3β sequence + peptide sequence; 2) TCR CDR3β sequence + antigen sequence + HLA sequence; 3) TCR CDR3β sequence + antigen sequence + HLA sequence + pHLA structure; 4) TCR CDR3αβ sequence + antigen sequence; 5) TCR CDR3αβ sequence + antigen sequence + HLA sequence; 6) TCR CDR3αβ sequence + antigen sequence + HLA sequence + pHLA structure. We conducted feature ablation tests to assess each combination’s contribution to antigen immunogenicity prediction. As depicted in Figure 2b, the results indicate that the sixth feature combination yields the most accurate predictions. Notably, omitting HLA data or TCR CDR3α information leads to a significant decline in model performance, underscoring the critical importance of incorporating both HLA and TCR CDR3α data for precise antigen immunogenicity prediction especially the structure information from pHLA.

### 3. Comparative Evaluation of MultiTAP with Existing Immunogenicity Prediction Models

#### 3.1. MultiTAP has better performance in independent tests

Based on the TPHD dataset, we compared MultiTAP with the widely recognized immunogenicity prediction tools NetTCR-2.0 and ERGO-I. We conducted independent testing to evaluate model performance, ensuring that issues of overfitting and underfitting were mitigated by using an independent test set. Training and test sets were derived from TPHD (Figure 3a), and algorithm training and evaluation were performed for MultiTAP, NetTCR-2.0, and ERGO-I.

**Fig. 3:**
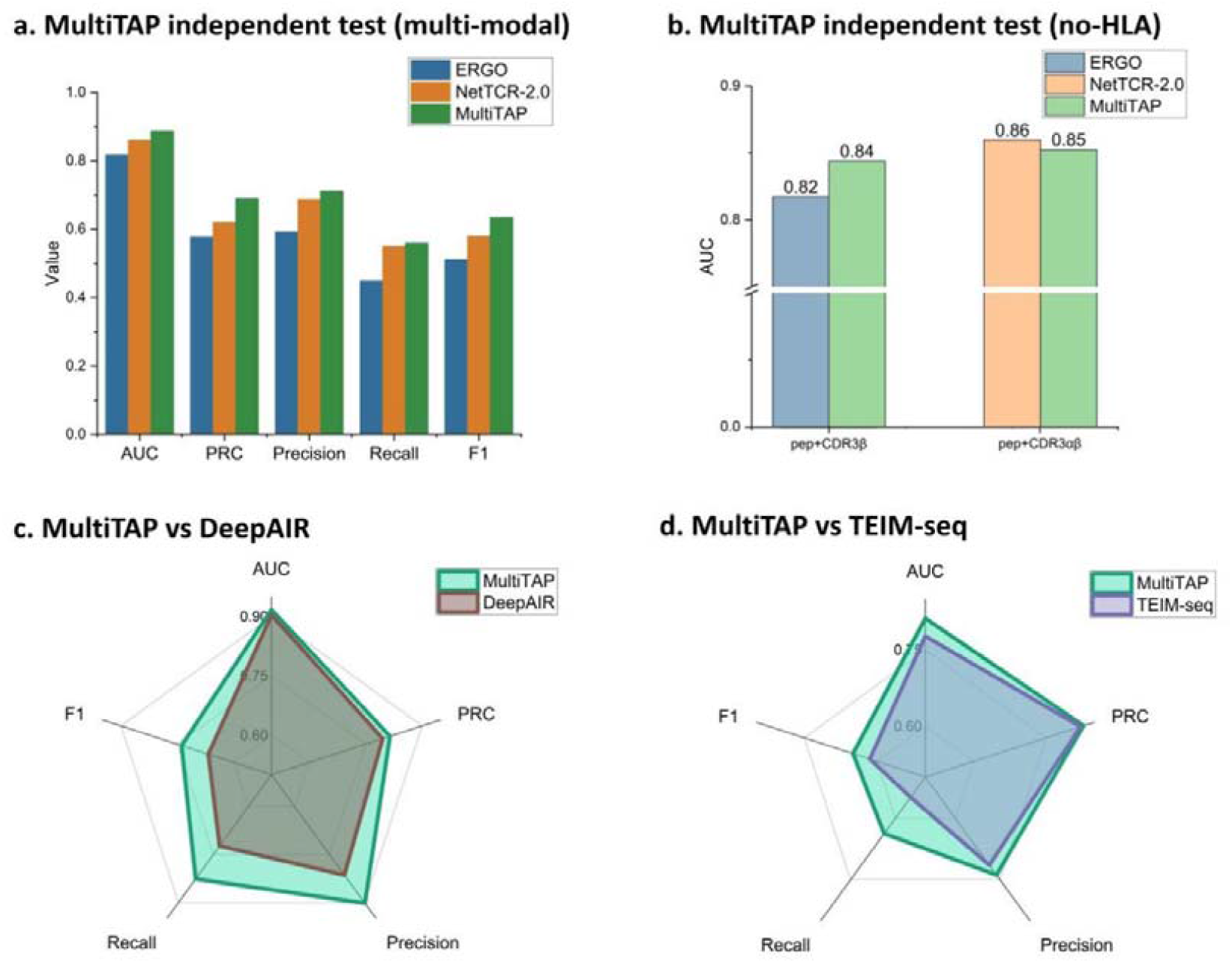
Comparison of predictive performance between MultiTAP and other algorithms. **a**, AUC, PRC, Precision, Recall and F1 scores for MultiTAP (multi-modal), ERGO and NetTCR-2.0 based on the same training set and the same test set from TPHD. b, Comparison between MultiTAP (no-HLA) and competing tools with the same feature types used for training. **c**, AUC, PRC, Precision, Recall and F1 scores for MultiTAP (multi-modal) and DeepAIR based on the same training set. d, AUC, PRC, Precision, Recall and F1 scores for MultiTAP (multi-modal) and TEIM-seq based on the same training set.

Performance was assessed using five standard binary classification metrics: AUC, PRC, Precision, Recall, and F1. The results indicated that, on the independent TPHD test set, NetTCR-2.0 and ERGO-I exhibited relatively lower performance (Figure 3b), whereas MultiTAP significantly outperformed both (Figure 3b).

It is notable that NetTCR-2.0 and ERGO-I do not incorporate HLA information, suggesting that MultiTAP’s enhanced performance may be attributed to its rich, multi-modal feature set. To confirm this, we evaluated model performance under comparable feature combinations. When aligning MultiTAP’s features with those used by NetTCR-2.0 and ERGO-I, MultiTAP’s performance notably decreased compared to its multi-modal configuration (Figure 3c), yet it matched or exceeded the performance of these models. This finding underscores the critical role of HLA information in accurately predicting antigens immunogenicity and demonstrates the advantage of multi-modal approaches in addressing such challenges.

We also noted that DeepAIR^49^ and TEIM^50^ excelled in predicting TCR-epitope interactions. However, because the TPHD dataset does not contain TCR-related genetic information (e.g., VDJ recombination), we could not conduct fair testing for DeepAIR and TEIM using TPHD. To address this, we trained and tested the MultiTAP model on the benchmark datasets used by DeepAIR and TEIM to facilitate an appropriate comparison (Figure 3d). The experimental outcomes revealed that MultiTAP continued to show superior predictive performance in independent tests.

### 3.2. MultiTAP achieved good performance in out-of distribution (OOD) test

To assess the generalization capability of the MultiTAP model, we conducted an out-of-distribution (OOD) test. OOD tests are essential in evaluating model stability and robustness, particularly by testing the model on datasets that differ from those used for training. This method minimizes the risk of artificially inflated performance due to similarities between training and test sets, which is vital in biomedical research where models must generalize across highly variable biological data to be reliable and practical for real-world applications^51,52^.

The need for robust OOD testing is particularly relevant when predicting the immunogenicity of tumor antigen epitopes. Clinical observations have shown that HLA phenotypes significantly restrict epitope presentation and immune activation, with substantial variations across populations.^53–55^. These differences limit the applicability of most current models to HLAs with significant diversity, presenting challenges for developing broad-spectrum tumor peptide vaccines. Additionally, the range of epitopes produced by various tumor types is vast^56,57^ and subject to issues like tumor immune evasion, leading to a dynamically shifting antigen landscape^58,59^. To address these challenges, we introduced an innovative clustered sequence-OOD sampling method to partition training and test data for OOD tests on TPHD. This method emphasizes sequence-level differences and clusters sequence data using multiple sequence alignment before executing a “hard split “^60^, achieving maximal differentiation between training and test sets based on sequence classification.

Utilizing sequence-OOD sampling, we conducted HLA-OOD and Antigen-OOD tests to assess MultiTAP’s performance in the context of HLA phenotype variability and the dynamic nature of epitope distribution. Figures 4a and 4b show the results of these tests, where MultiTAP with multi-modal conditions was compared against NetTCR-2.0 and ERGO. With an AUC threshold of 0.5, both NetTCR-2.0 and ERGO exhibited marked declines in predictive capability, with AUC values approaching 0.5, indicating poor performance in OOD scenarios. Conversely, MultiTAP maintained superior predictive performance, demonstrating its robustness in classifying antigens in diverse data settings. This highlights MultiTAP’s enhanced generalization across different HLA phenotypes and epitope sequences, showcasing its potential as a reliable tool for cancer vaccine and immunotherapy development.

**Fig. 4:**
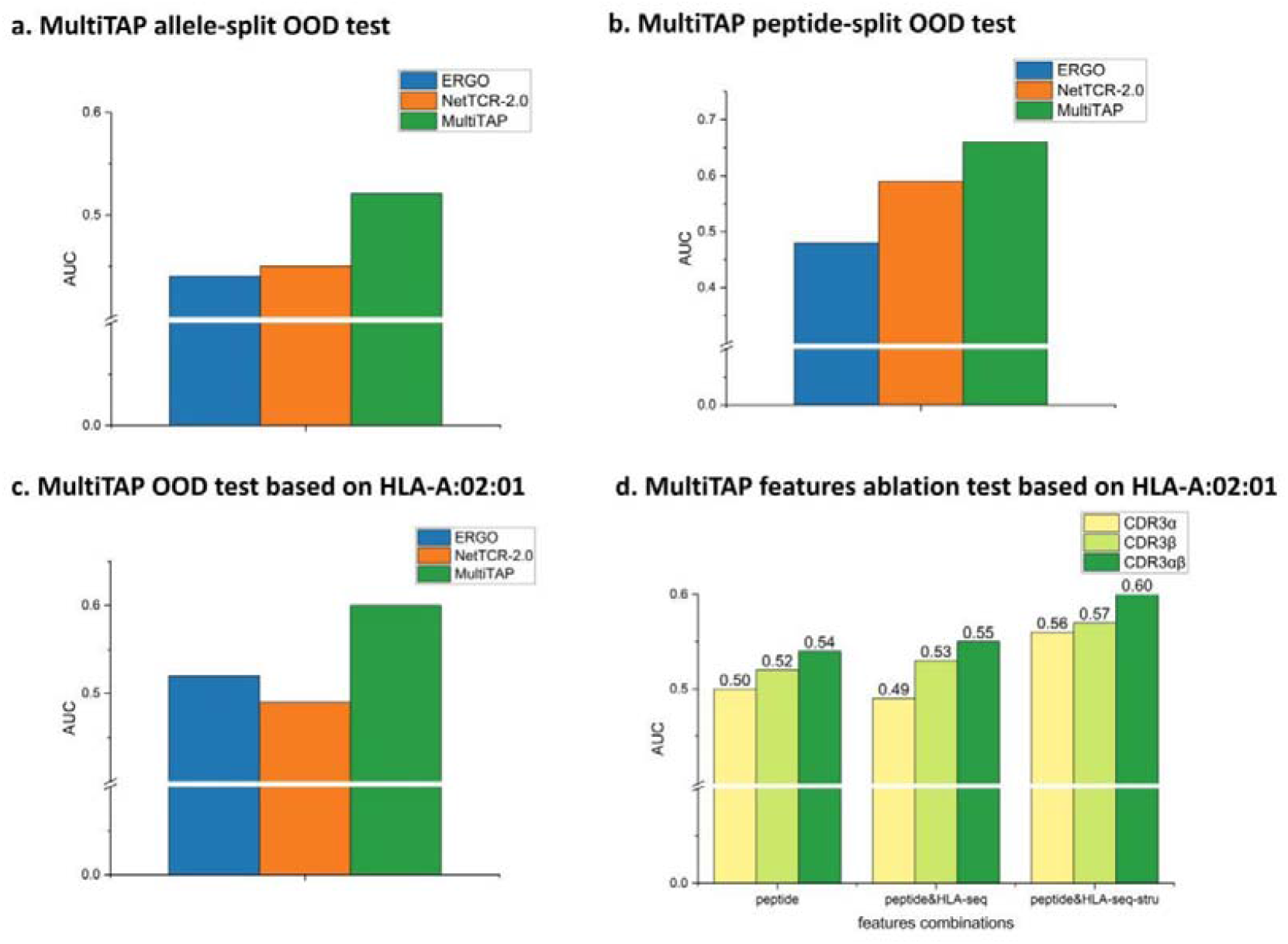
Results of MultiTAP and competing tools in allele-split and peptide-split out-of distribution (OOD) tests. **a**, Predictive accuracy of MultiTAP, ERGO, and NetTCR-2.0 in allele-split OOD tests, demonstrating the robustness of MultiTAP in handling unseen HLA alleles. **b**, Accuracy comparison of MultiTAP, ERGO, and NetTCR-2.0 in peptide-split OOD tests, highlighting MultiTAP’s superior performance with novel antigen sequences. **c**, Evaluation of predictive accuracy when using HLA-A:02:01 as the test set phenotype, with training conducted on other phenotypes, showing MultiTAP’s advantage over ERGO and NetTCR-2.0 in cross-phenotype generalization. **d**, Results from feature ablation experiments performed on the training and test sets from **c**, demonstrating the contribution of specific features (e.g., CDR3α sequence, HLA sequence, HLA structure) to MultiTAP’s overall predictive performance.

We also applied a “hard split” approach, training on TPHD data from the HLA-A:02:01 phenotype and testing on other phenotypes. MultiTAP’s stability again exceeded that of NetTCR-2.0 and ERGO-I (Figure 4c). Notably, we observed the positive impact of incorporating HLA structural information on MultiTAP’s performance stability (Figure 4d). These findings further validate that the multi-modal architecture of MultiTAP, enriched with diverse data types, not only enhances predictive accuracy but also bolsters the model’s robustness across a broader range of HLA phenotypes and antigenic variability.

### 4. MultiTAP uncovers the underlying patterns of TAP binding

In the MultiTAP model structure, we utilized an attention mechanism to interactively map the sequence and structural information of pHLA with TCR and epitope, thereby obtaining a complex integration state of information about the epitope within the “pore structure” formed by the TCR-HLA complex. Here, we observed the complex integration state of information about epitopes generated by the model knowledge. In several explainable tools, attention has been used to represent the importance of features^49,50^. In this context, we associated the generated pHLA with the final output and, through the weight allocation mechanism of attention, observed the weights of different residues in the complex information integration state of pHLA.

This approach allowed us to identify which residues and structural elements of the pHLA complex play a critical role in the interaction with TCR and the presentation of epitopes. By analyzing the attention weights, we could pinpoint specific areas within the complex that are crucial for epitope recognition and TCR binding. This insight is invaluable for understanding the mechanisms of immune recognition and has implications for designing more effective cancer vaccines and immunotherapies by highlighting which aspects of the TCR-peptide-HLA interaction are most influential in determining immunogenicity.

Initially, we observed the relationship between the complex integration state of pHLA information for epitopes of different lengths and the immunogenicity of antigenic peptides. Without considering HLA classification (Figure 5a), we found that MultiTAP, when predicting epitope immunogenicity, tends to focus attention on the 6th, 7th, 1st, and 2nd residues of the epitope. Structurally, the focused residues vary; some have carbon atoms at average distances farther from HLA, while others concentrate on conserved sites of the non-covalent relationship between HLA and the epitope. This indicates that the complex integration state information of epitopes generated by model knowledge includes an understanding of the structural information of the ternary complex.

**Fig. 5:**
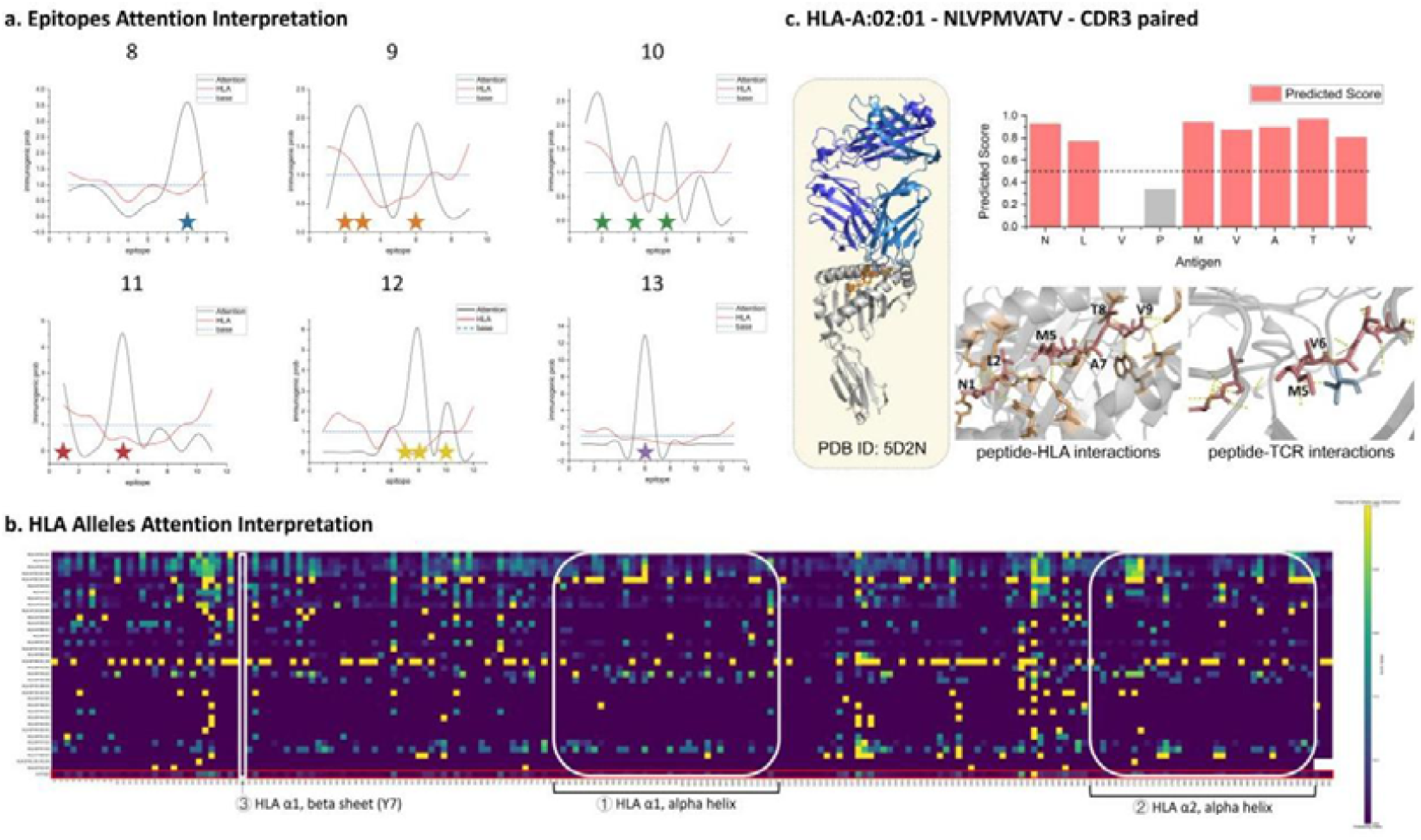
Analysis of MultiTAP’s interpretation and integration of pHLA information in immunogenicity prediction. **a**, Attention distribution across epitope residues of varying lengths, highlighting that MultiTAP focuses on the 6th, 7th, 1st, and 2nd residues when predicting immunogenicity, irrespective of HLA classification. The structural analysis reveals that some targeted residues have carbon atoms positioned at greater average distances from HLA, while others coincide with conserved non-covalent interaction sites between HLA and the epitope, indicating the model’s understanding of ternary complex structural information. **b**, Evaluation of attention weights across different HLA phenotypes, showing that MultiTAP does not directly encode epitope-HLA interactions based solely on HLA sequences. **c**, Analysis of positive samples with crystal-resolved TCR-epitope-HLA complexes, demonstrating that MultiTAP identifies non-covalent interactions between epitope residues and both TCR and HLA. This supports the notion that MultiTAP indirectly learns HLA-related information, integrating this knowledge into its epitope representation and reflecting the biological importance of HLA in the T-cell immune response.

Subsequently, we analyzed the value of different HLA combinations in the complex integration state. By observing the weight information of amino acid sites on different HLA phenotypes, we found that MultiTAP did not directly observe the epitope-HLA interaction on the HLA sequence (Figure 5b). Further, upon examining positive sample data with resolved crystal structures of the TCR-epitope-HLA ternary complex, we discovered that MultiTAP noticed the non-covalent structural interactions between the epitope residues and both TCR and HLA (Figure 5c). This suggests that MultiTAP’s learning of HLA information is indirect, with the role of HLA phenotype in predicting immunogenicity integrated into MultiTAP’s learning about epitopes. This aligns with the upstream and downstream signaling relationships in the T-cell immune process and reaffirms the significant value of HLA information in predicting the immunogenicity of epitopes.

### 5. MultiTAP has good performance for HLA-A:11:01 allele antigens prediction

Currently, the HLA allele in the vast majority of public datasets, predominantly HLA-A:02:01, has the highest proportion. However, there is still insufficient evidence to indicate that the HLA-A:02:01 phenotype is widely distributed in the East Asian region. HLA phenotype restriction means that we also need to focus on the immunogenicity after epitope presentation of HLA phenotypes that are broadly distributed in the East Asian region and employ suitable tools to assist in the screening and design of such types of antigens.

According to literature reports, the predominant HLA allele in common East Asian ethnicities is HLA-A:11:01^61,62^. In this context, we have collected TAP-positive sample data of East Asian hepatitis B patients from some literature (Methods-dataset-HLA-A:11:01 dataset). The collected HLA-A:11:01 dataset mainly includes four antigens, which are STLPETAVVRR, RSQSPRRRRSK, LVVDFSQFSR, and KTAYSHLSTSK. We have created the binary complex crystal structures of these four antigens with HLA-A:11:01 using xTrimo-Multimer (Figure 6a).

**Fig. 6:**
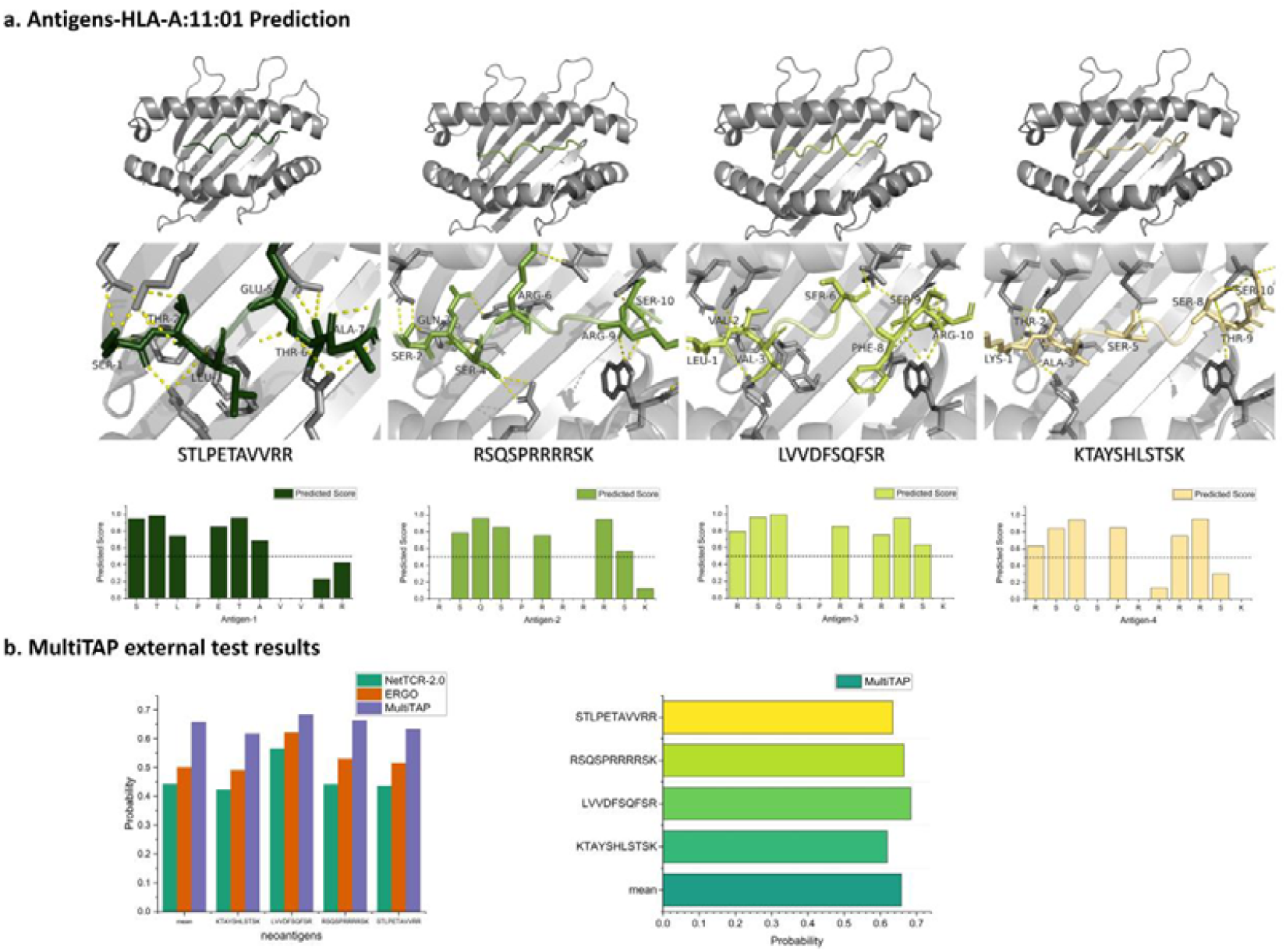
Performance analysis of MultiTAP on the HLA-A:11:01 dataset of East Asian hepatitis B patients. **a**, Crystal structures of the binary complexes formed by four antigens with HLA-A:11:01: STLPETAVVRR, RSQSPRRRRSK, LVVDFSQFSR, and KTAYSHLSTSK (structures are generated by the xTrimo-Multimer prediction model). **b**, Comparison of immunogenicity prediction performance for these antigens using the MultiTAP model trained on the TPHD dataset. The results demonstrate that MultiTAP achieved an average prediction AUC exceeding 0.6 across all four antigens, with a particularly notable AUC close to 0.7 for LVVDFSQFSR, indicating the model’s enhanced predictive capability for antigen binding and immunogenicity in the HLA-A:11:01 context.

Based on the MultiTAP model trained with TPHD, we observed the performance of immunogenicity prediction after the presentation of antigens based on HLA-A:11:01. Across these four antigens, the average prediction AUC of MultiTAP exceeded 0.6, indicating a stronger capability of MultiTAP in predicting the immunogenicity binding of antigens on the HLA-A:11:01 phenotype compared to other models (Figure 6b). Notably, the immunogenicity accuracy for LVVDFSQFSR reached an AUC close to 0.7 (Figure 6b).

## Discussion

In this study, we propose MultiTAP, a multi-modal approach that achieves better epitope immunogenicity prediction under different dataset partition settings for precision screening of immunogenic tumor antigens. On a constructed dataset namely TPHD that innovatively integrates sequence with the structures of pHLA, MultiTAP demonstrates that a multi-modal deep learning architecture can effectively improve prediction of immunogenicity for tumor antigens. This heralds a leap forward in the precision tumor antigens screening and the development of cancer immunotherapies.

MultiTAP demonstrates that the addition of HLA information enhances the prediction of epitope immunogenicity, especially when sequence and structural information are used together, offering new perspectives for future epitope immunogenicity model feature inputs. Compared to multiple baselines, MultiTAP exhibited better performance in accurate prediction of epitope immunogenicity. Unlike most models that use a “hard split”, MultiTAP innovatively employs sequence-OOD sampling to test generalization ability at an out-of-distribution test set, showcasing MultiTAP’s excellent capability in mining immunogenic antigen peptides on a real clinical dataset characterized by HLA-A:11:01 phenotype. Moreover, we find that the focus of epitope immunogenicity prediction still lies in the learning and utilization of epitope information, with algorithm optimization significantly impacting model predictive performance.

Due to the lack of crystal structure data, we resorted to using xTrimo-Multimer protein structure prediction model to create structural data; however, the limited accuracy and constraints might have hindered a deeper understanding of the TCR-peptide-HLA complex binding patterns by MultiTAP. Thus we hope to access more high-quality crystal structure data in future. Moving forward, MultiTAP will continue to delve into the binding patterns of TCR-peptide-HLA complexes, offering better suggestions for cancer immunotherapies that target tumor antigens.

## Methods

### 1. Design of MultiTAP

The MultiTAP algorithm employs a multi-modal algorithmic structure, utilizing features from three types of protein sequence data and one type of protein structural data for model training and testing.

CDR3-paired TCR protein sequence features are extracted from an unsupervised pre-trained protein language model, ESM-1b, producing feature vectors *Seq*_*TCR*_ of a specified size (*batch, CDR3_sequence_length_max*, 1280). Peptide-like antigen sequence features are extracted using one-hot encoding, classifying amino acids in the protein sequence into 20 natural amino acid categories and one unknown amino acid category, resulting in feature vectors *Seq*_*antigen*_ of a certain size *(*batch, peptide_sequence_length_max, 21*)*. HLA protein sequence features are extracted using a method based on multiple Position Specific Scoring Matrix (PSSM) alignments, utilizing NCBI Psi-Blast, producing feature vectors *Seq*_*HLA*_ of a specified size *(*batch, HLA_sequence_length_max, 20*)*. For structural data feature extraction, based on pHLA binary protein complex structure PDB data, we extract the Laplacian matrix and adjacency matrix *Stru*_AHLA_ using residues as nodes and their spatial distances as edges, which are then further input into MultiTAP.

MultiTAP’s first layer of embedding processes features *Seq*_TCR_, *Seq*_*Antigen*_, *Seq*_*HLA*_ and *Stru*_*AHLA*_ respectively, after which feature transformations are applied to produce transformed features *Embedding*_*TCR*_, *Embedding*_*Antigen*_, *Embedding*_*HLA*_:

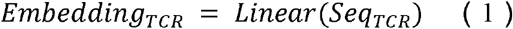

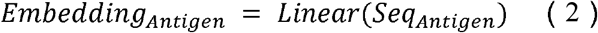

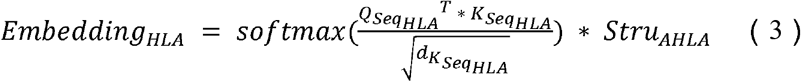

*Embedding*_*HLA*_ *is* obtained by joint representation from *Stru*_*AHLA*_ and the self-attention matrix of the extracted features *Seq*_*HLA*_.

The second layer of MultiTAP consists of a three-layer gated convolutional neural network (GCNN) processing *Embedding*_*TCR*_, and a three-layer attention mechanism handling interactions between *Embedding*_*Antigen*_ and *Embedding*_*HLA*_. The kernel size of the GCNN is 3, described by the formula:

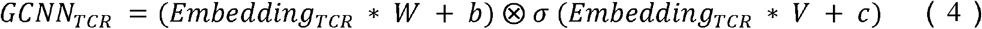

Where the parameters *W, V, b, c* are learned. GCNNs are widely used in NLP for parallel and efficient processing of complex sequence information^63^. After passing through the three-layer GCNN, the transformed features *C* _*TCR*_ are obtained.

The attention mechanism formula is as follows (5) and (6), facilitating interaction between *Embedding*_*Antigen*_ and *Embedding*_*HLA*_, projecting HLA information onto peptide sequence information based on residue positions. After the three-layer attention mechanism, the feature vector *Attention*_*AHLA*_ is obtained.

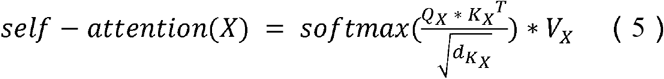

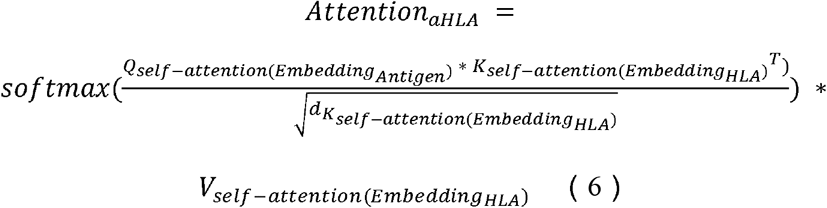

The third layer of MultiTAP involves three layers of multi-head attention for processing interactions between TCR and HLA-peptide features, with the calculation formulas for the multi-head attention mechanism provided. *Multiheads - attention*_*TCR-AHLA*_ utilizes the multi-head attention mechanism to project TCR information onto the pHLA, thereby predicting the immunogenic capacity of the peptide-based on pHLA.

Finally, MultiTAP utilizes two linear fully connected layers as the label predicting layer, producing model prediction results of (*batch, 2*) to determine the immunogenic potential of peptides.

## 2. Dataset

### 2.1. TPHD: TCR-peptide-HLA Dataset

The TCR-peptide-HLA Dataset is designed for multi-modal peptide immunogenicity prediction modeling. Our dataset integrates data from three public databases: IEDB, McPAS-TCR, and VDJdb. To validate the significance of CDR3-paired data, HLA sequence information, and HLA structural information in the TPHD dataset, we selectively compiled positive sample data that included CDR3-paired sequences, HLA sequences/HLA phenotype names, and peptide sequences from these databases. After collecting data from the three databases, we cleaned and deduplicated the dataset to ensure there were no repeated CDR3β-HLA-peptide positive samples.

We used the xTrimo-Multimer model to create HLA structural data. Since our MultiTAP algorithm aims to predict peptide immunogenicity, we focused on addressing the interaction problem between TCR and pHLA. Therefore, we did not create HLA structural data in isolation but generated pHLA binary complex structural data instead. We created 1017 different pHLA binary complex structures, covering all the pHLA requirements in the entire dataset of TCR-pHLA combinations.

Subsequently, we randomized the combinations of TCR and pHLA to generate negative samples fivefold. We also conducted a data correlation analysis between the sequence information of positive and negative samples and their immunogenicity outcomes, validating the feasibility of this negative sample generation method.

### 2.2. HLA-A:11:01 Dataset

To validate the generalization capability of the MultiTAP model, we designed a dataset exclusively comprising the HLA-A:11:01 phenotype. The HLA-A:11:01 phenotype is considered to be widely distributed among East Asian populations, accounting for approximately 12.9% of the TPHD dataset. A dataset based on the HLA-A:11:01 phenotype is sufficiently representative to verify the model’s generalization ability.

This dataset’s data originates from the work of Cheng et al.^64^, which documented the immunogenicity of four types of antigen peptides in hepatitis B patients. Based on this literature, we obtained 4701 TCR-pHLA data entries, thereby creating the HLA-A:11:01 dataset.

### 2.3. Benchmark Datasets

In independent testing, we also conducted algorithm comparisons based on datasets from other benchmarks. Since most algorithms have not yet utilized both CDR3-paired data and HLA data for model training and testing, we chose the datasets used by DeepAIR and TEIM for independent testing comparisons.

DeepAIR introduced various methods for antigen-receptor interaction prediction. The dataset related to TCR-antigen has been made public on GitHub: https://github.com/TencentAILabHealthcare/DeepAIR. DeepAIR utilizes TCR-peptide-HLA data for model construction, allowing us to obtain protein sequence information related to TCR-peptide-HLA from their public dataset and create pHLA structural information using xTrimo-Multimer.

TEIM is a tool that uses protein sequence information for peptide immunogenicity prediction. Its model input includes TCR protein sequences, antigen protein sequences, and HLA protein sequences. The public dataset GitHub link is: https://github.com/pengxingang/TEIM. Here, we also created pHLA structural data using xTrimo-Multimer.

Subsequently, we used the modified DeepAIR dataset and the modified TEIM dataset for training MultiTAP, comparing the antigen peptide immunogenicity prediction capabilities between MultiTAP and DeepAIR, and between MultiTAP and

TEIM.

## 3. Sampling Methods

### 3.1. Balanced sampling

The balanced sampling method was utilized for dividing the training and testing datasets in the independent testing of MultiTAP. Based on sequence information and sample labels, we balanced the samples in both the training and testing datasets. We first meticulously categorized the TPHD dataset from three aspects: label categories, HLA phenotype types, and peptide categories. Then, we classified all the data accordingly and subsequently sampled training and testing data at an 8:2 ratio for each category.

This method ensured that the proportions of positive and negative samples, HLA phenotype ratios, and peptide category ratios were consistent between the training and testing datasets, creating training and testing sets with uniform data distributions.

### 3.2. Sequence-OOD sampling

The sequence-OOD (Out-Of-Distribution) sampling method was used for dividing the training and testing datasets in the OOD testing of MultiTAP. The sequence-OOD sampling approach primarily consists of two steps: category setting and sequence similarity clustering.

Firstly, the category-setting method consistent with balanced sampling was employed. The TPHD dataset was meticulously categorized from three perspectives: label categories, HLA phenotype types, and peptide categories, with all the data classified accordingly. Then, using the k-means clustering method based on protein sequence information, we clustered the data for a variable for which we needed to design the OOD experiment.

In this study, we conducted experiments for both HLA-OOD and peptide-OOD. Therefore, we performed k-means clustering of protein sequence information separately for the HLA variable and the peptide variable. Based on the clustering results, we divided the training and testing datasets for the OOD experiments in an 8:2 ratio. Detailed clustering results and proportion divisions can be found on GitHub (https://github.com/ruofanjin/MultiTAP).

## 4. Baselines

For our experiments, we primarily used two common baselines: ERGO-I and NetTCR-2.0. Both methods have been used as baselines in various articles^21,65^.

ERGO-I predicts peptide-TCR binding using either long-short term memory (LSTM) networks or autoencoders (AE). Both network architectures were trained on datasets derived from VDJdb and/or McPAS. In this case, we used ERGO-AE as one of our baselines, based on the official code provided (https://github.com/louzounlab/ERGO), we retrained ERGO-I on the TPHD dataset from scratch and compared it with MultiTAP in independent and OOD tests.

NetTCR-2.0 is a linear neural network model trained on data from The Immune Epitope Database (IEDB) and VDJdb. Here, based on the CDR3-paired and peptide sequence data in the TPHD dataset, we retrained NetTCR-2.0 (https://github.com/mnielLab/NetTCR-2.0) from scratch and compared it with MultiTAP in independent and OOD tests.

## 5. Experiments

To fully explore the feasibility and application value of MultiTAP, this study conducted independent tests, HLA-OOD tests, peptide-OOD tests, and external tests.

In the independent test, we employed the balanced sampling method to divide the training and testing datasets based on the TPHD dataset (https://github.com/ruofanjin/MultiTAP/data/TPHD). Here, we conducted independent testing experiments comparing MultiTAP, ERGO-I, and NetTCR-2.0, using five evaluation metrics for assessing model prediction capabilities: AUC, PRC, precision, recall, and F1 score.

For the HLA-OOD test and peptide-OOD test, we used the sequence-OOD sampling method to divide the training and testing sets for the HLA and peptide variables, respectively.

In the external test, we tested the MultiTAP model, which was trained on the TPHD dataset, on the HLA-A:11:01 dataset. Since the HLA-A:11:01 dataset consists entirely of positive samples, we only evaluated the accuracy of MultiTAP’s predictions for positive samples.

## Supporting information

Cover Letter - Only for Editorial

## Data availability

The training and testing datasets are shared on our GitHub repository: https://github.com/ruofanjin/MultiTAP. The raw data is from IEDB, McPAS-TCR, and VDJdb can also be found on our Git Hub (https://github.com/ruofanjin/MultiTAP/data/raw_data).

## Code availability

The source codes and model weights of MultiTAP are available on GitHub (https://github.com/ruofanjin/MultiTAP).

## Acknowledgements

We thank Qing Ye, Kevin Chun Chan and Dong Zhang for helpful discussions. This work was partially supported by the National Key R&D Program of China (2021YFF1200404 and 2021YFA1201200), the National Natural Science Foundation of China (U1967217), the National Center of Technology Innovation for Biopharmaceuticals (NCTIB2022HS02010), Shanghai Artificial Intelligence Lab (P22KN00272), the National Independent Innovation Demonstration Zone Shanghai Zhangjiang Major Projects (ZJZX2020014), the Starry Night Science Fund of Zhejiang University Shanghai Institute for Advanced Study (SN-ZJU-SIAS-003) and Zhejiang University Global Partnership Fund (188170+194452409/004).

## Contributions

R.Z. conceived the idea and designed the research. R.Z., T.H. and K.H. provided in silico experiments’ guidance for the research. R.J. designed the multi-modal deep learning framework MultiTAP and wrote the software with the support from J.G. and G.Z.. R.J. performed structure modeling of pHLA complexes. R.J. and R.Z. wrote the manuscript. All authors participated in discussions and revisions of the manuscript.

