## Supplementary material for "A Multi-Modal Deep Learning Framework with Both Sequence and Structure for Tumor Antigens Prediction": Cover Letter - Only for Editorial

ZHEJIANG UNIVERSITY

Qiushi Chair Professor and Director
Institute of Quantitative Biology
Zhejiang University, China


Nov 5th, 2024

Dear editors,

We are pleased to submit the manuscript titled "**A Multi-Modal Deep Learning Framework with Both Sequence and Structure for Tumor Antigens Prediction**" for your kind consideration of publication in *BioRxiv*. The following is a brief description of our work and its significance in the future development of neoantigen-based personalized immune-therapies.

Tumor antigens have emerged as pivotal targets in cancer immunotherapy, garnering significant interest within the scientific community and becoming a focal point of extensive research efforts. These antigens are presented on the cell surface by MHC class I molecules in the tumor microenvironment and subsequently recognized by CD8+ T cells, initiating tumor-specific immune responses and facilitating T cell-mediated tumor cell elimination. Originating primarily from tumor-specific somatic mutations and absent in normal tissues, these antigens exhibit unique immunogenicity that effectively activates the immune system and elicits robust T cell responses. Immunotherapeutic strategies targeting these neoantigens have shown promise in augmenting the antitumor efficacy of existing treatments, particularly in the development and implementation of personalized therapeutic approaches.

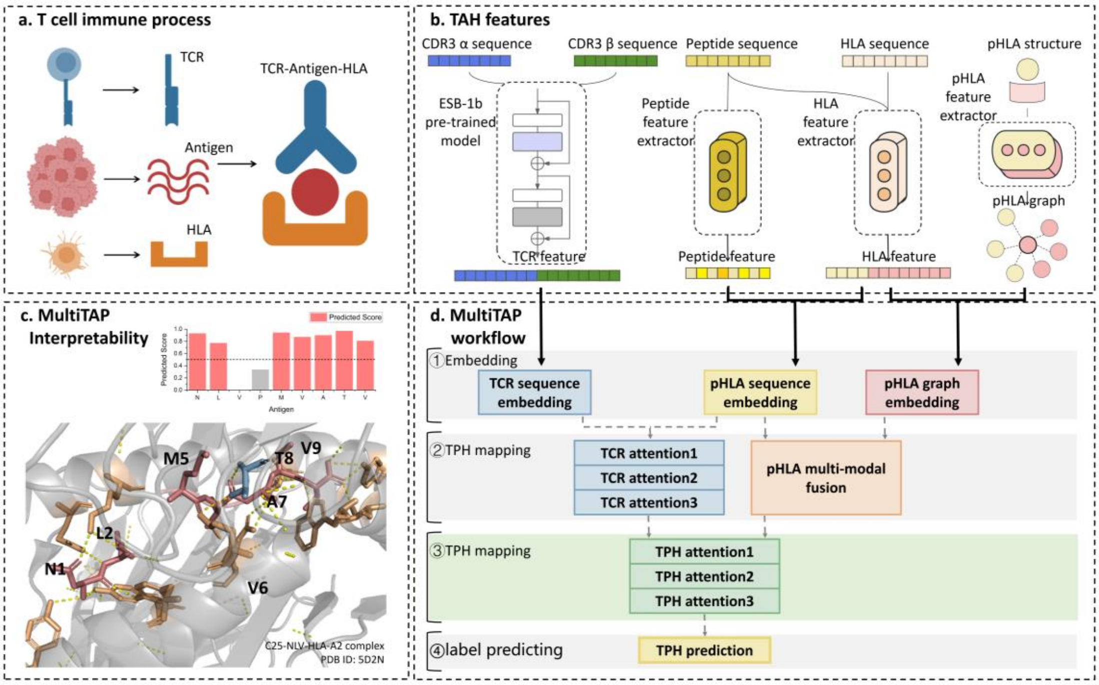


**Figure R1**. Overall architecture of the MutiTAP model.

However, precision screening of immunogenic tumor antigens remains a great challenge due to human leukocyte antigen (HLA) restriction and tumor antigen escape. Here, we introduce MultiTAP (Multi-modal Tumor Antigen Predictor; see **Figure R1**), a pioneering multi-modal framework with HLA structure incorporating an attention mechanism designed to accurately identify tumor antigens with immunogenic properties. By constructing the multi-modal TCR-peptide-HLA Dataset (TPHD) and integrating its sequence and structure, we perform antigen feature enhancement using peptide-HLA (pHLA) structural features at the residue level, achieving interpretable prediction of immunogenicity for tumor antigens. Relative to existing baseline models, the MultiTAP framework has exhibited superior efficacy in predicting the immunogenicity of tumor antigens. Through comprehensive out-of-distribution (OOD) assessments, MultiTAP has maintained predictive robustness across diverse HLA phenotypes and the continuously evolving landscape of epitope distributions. Overall, MultiTAP presents a brand-new and promising approach for cancer immunotherapies that target tumor antigens.

To conclude, we successfully provide a multi-modal deep learning framework, MultiTAP, to enhance the efficacy of neoantigen prediction for future potential personalized immunotherapy. We believe that this work would have broad impacts for the community.

We are extremely grateful for your time and kind consideration in evaluating this work and we look forward to hearing from you in the near future.

Thanks.

Sincerely,

Ruhong Zhou, PhD
Zhejiang Univ and Columbia Univ
