## Supplementary material for "A Multi-Modal Deep Learning Framework with Both Sequence and Structure for Tumor Antigens Prediction": Tables and Figures

### Contents


1 Supplymentary Tables

1.1 Results of features ablation tests of MultiTAP

1.2 Results of independent tests of MultiTAP

1.3 Results of out-of distribution tests of MultiTAP

1.4 Results of hard split tests of MultiTAP

1.5 Results of external test of MultiTAP

2 Comparison of TCR-pHLA Structure Prediction Tools

2.1 Introduction

2.2 Prediction accuracy of TCR-pHLA crystal structure data

2.3 Time consumption on TCR-pHLA structure prediction

3 Methods for Negative Samples Generation

4 Interpretability of MultiTAP

4.1 Interpretability based on peptide amino acid sites of different lengths

4.2 Interpretability based on HLA amino acid sites of different alleles

### 1 Supplymentary Tables

#### 1.1 Results of features ablation tests of MultiTAP

Table S1: **Results of features ablation tests of MultiTAP**

| | peptide | peptide & HLA-seq | Peptide & HLA-seq & HLA-stru |
|---|---|---|---|
| CDR3α | 0.79707 | 0.81933 | 0.83216 |
| CDR3β | 0.84388 | 0.86572 | 0.88096 |
| CDR3αβ | 0.85244 | 0.87592 | 0.88728 |

#### 1.2 Results of independent tests of MultiTAP

Table S2: **Results of independent tests of MultiTAP based on TPHD**

| | AUC | PRC | Precision | Recall | F1 |
|---|---|---|---|---|---|
| MultiTAP | 0.887 | 0.690 | 0.711 | 0.560 | 0.635 |
| NetTCR-2.0 | 0.860 | 0.620 | 0.687 | 0.549 | 0.579 |
| ERGO | 0.817 | 0.577 | 0.592 | 0.449 | 0.511 |

Table S3: **Results of independent tests of MultiTAP based on DeepAIR dataset**

| | AUC | PRC | Precision | Recall | F1 |
|---|---|---|---|---|---|
| MultiTAP | 0.917 | 0.815 | 0.902 | 0.826 | 0.739 |
| DeepAIR | 0.904 | 0.796 | 0.814 | 0.723 | 0.668 |

Table S4: **Results of independent tests of MultiTAP based on TEIM-seq dataset**

| | AUC | PRC | Precision | Recall | F1 |
|---|---|---|---|---|---|
| MultiTAP | 0.812 | 0.827 | 0.738 | 0.638 | 0.649 |
| TEIM-seq | 0.777 | 0.821 | 0.714 | 0.549 | 0.615 |

#### 1.3 Results of out-of distribution tests of MultiTAP

Table S5: **Results of HLA allele out-of distribution tests of MultiTAP**

| | ERGO | NetTCR-2.0 | MultiTAP |
|---|---|---|---|
| AUC | 0.44 | 0.45 | 0.521 |

Table S6: **Results of peptide out-of distribution tests of MultiTAP**

| | ERGO | NetTCR-2.0 | MultiTAP |
|---|---|---|---|
| AUC | 0.48 | 0.59 | 0.66 |

#### 1.4 Results of hard split tests of MultiTAP

Table S7: **Results of MultiTAP based on HLA-A:02:01 allele data for training**

| | ERGO | NetTCR-2.0 | MultiTAP |
|---|---|---|---|
| AUC | 0.52 | 0.49 | 0.6 |

#### 1.5 Results of external test of MultiTAP

Table S8: **Results of external test of MultiTAP**

| | ERGO | NetTCR-2.0 | MultiTAP |
|---|---|---|---|
| KTAYSHLSTSK | 0.491 | 0.424 | 0.619 |
| LVVDFSQFSR | 0.622 | 0.567 | 0.684 |
| RSQSPRRRRSK | 0.531 | 0.442 | 0.665 |
| STLPETAVVRR | 0.517 | 0.437 | 0.634 |

### 2 Comparison of TCR-pHLA Structure Prediction Tools

#### 2.1 Introduction

Alpha-Multimer is an advanced computational tool tailored for analyzing and predicting protein multimer interactions. Utilizing cutting-edge algorithms and machine learning, it models molecular interactions accurately, aiding in biomedical research and drug development. This software features a user-friendly interface and powerful visualization tools, making it essential for understanding complex protein behaviors and the impact of mutations on protein functionality in various biological contexts.

In this study, we conducted a model comparison among AlphaFold-Multimer and xTrimo-Multimer. Based on the TPHD dataset, which includes 131 entries with actual TCR-pHLA ternary complex crystal structures, we extracted the peptide-HLA parts and obtained 96 different peptide-HLA structures. Subsequently, using the sequence information from the PDB files, we generated structures from scratch for these 96 peptide-HLA structures using two different methods and scored the predictions using the RMSD metric. We also compared the time consumption for predicting the 96 peptide-HLA structures using these two methods. Combining prediction accuracy and time efficiency, we selected the peptide-HLA structures predicted by xTrimo-Multimer as the structural data for the TPHD dataset.

#### 2.2 Prediction accuracy of TCR-pHLA crystal structure data

Table S9: **RMSD of TCR-pHLA crystal structure data**

| | AlphaFold-Multimer_v3 | xTrimo-Multimer |
|---|---|---|
| average | 0.983 | 1.732 |

Table S10: **RMSD of peptide crystal structure data**

| | AlphaFold-Multimer_v3 | xTrimo-Multimer |
|---|---|---|
| average | **0.765** | **0.806** |

#### 2.3 Time consumption for TCR-pHLA complexes structure prediction

Table S10: **Time consumption for TCR-pHLA complexes structure prediction**

| | AlphaFold-Multimer_v3 | xTrimo-Multimer |
|---|---|---|
| average | 3163.29s | **137.18s** |

### 3 Methods for Negative Samples Generation

Generating negative samples is a common step in processing datasets for deep learning. In our dataset, as it consists solely of positive samples and lacks negative data, it is necessary for us to generate negative samples. A typical method for generating such samples involves random combinations based on our label categories. NetTCR-2.0 uses a method of generating negative samples randomly at five times the rate of positive samples for dataset creation, while ERGO employs a method of generating negative samples at ten times the rate for its dataset creation.

Here, we adopted a method of random recombination based on pHLA and TCR to generate negative samples. We believe that an effective negative sample generation method should meet the following criteria: 1) minimal false negatives; 2) it should not affect the model's description of the established true TCR-pHLA binding patterns; 3) there should be a discernible difference between positive and negative samples. To this end, we observed the positive correlation of positive and negative samples based on TPHD.

The pHLA attention results show the TCR's focus on each residue site of the pHLA sequence. We examined the pHLA attention results for both positive and negative samples in independent tests, as shown in Figure S1, where the x-axis represents the position of each residue site, and the y-axis shows the pHLA attention score's classification result at -1 or 1. We statistically compared the pHLA attention predictions for positive samples with those for negative samples. If the probability of a site being -1 or 1 exceeds 50% of the dataset, it is classified as -1 or 1; otherwise, it is 0. This produced two sets of data termed one_importance and zero_importance.

The correlation analysis between the two datasets, one_importance and zero_importance, yielded an R value of 0.111 and a p-value of 0.03. This indicates that there is no positive correlation between the positive and negative samples predicted by MultiTAP and that there is a significant difference between them. This validates our method for generating negative samples as both correct and feasible.

Figure S1: **Assessment of the Rationality of Negative Sample Generation Methods**

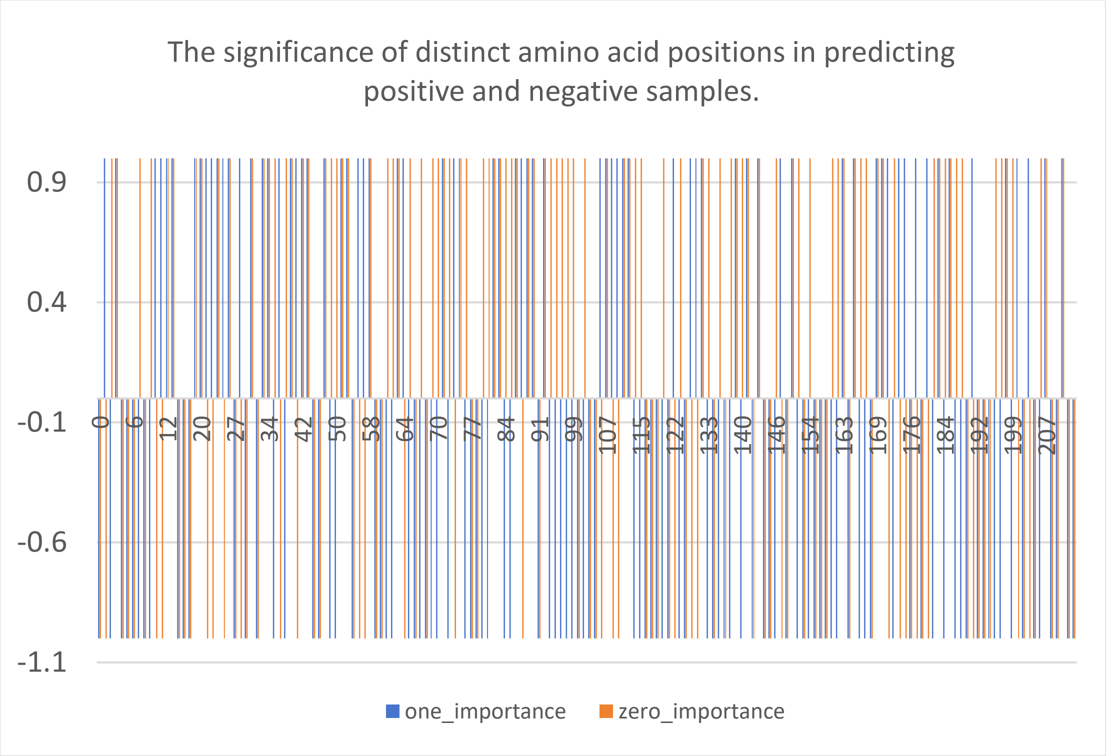

### 4 Interpretability of MultiTAP

#### 4.1 Interpretability based on peptide amino acid sites of different lengths

Table S11:

| | TCR | HLA |
|---|---|---|
| 1 | **7** | **1. 68** |
| 2 | 0 | **1. 54** |
| 3 | 0 | 0. 98 |
| 4 | 0 | 0. 42 |
| 5 | 0 | 0. 98 |
| 6 | 0 | 0. 84 |
| 7 | 0 | 0. 56 |

Table S12:

| | TCR | HLA |
|---|---|---|
| 1 | 0.8 | **1.393728223** |
| 2 | **1** | **1.142857143** |
| 3 | 0.6 | **1.031358885** |
| 4 | 0 | 0.473867596 |
| 5 | 0.4 | 0.947735192 |
| 6 | **1.2** | 0.808362369 |
| 7 | **3.6** | 0.780487805 |
| 8 | 0.4 | **1.421602787** |

Table S13:

| | TCR | HLA |
|---|---|---|
| 1 | 0.414713015 | **1.499750167** |
| 2 | **1.746160065** | **1.359593604** |
| 3 | **2.153597413** | 0.8994004 |
| 4 | 0.851253032 | 0.442205197 |
| 5 | 0.407437348 | 0.574117255 |
| 6 | **1.891673403** | 0.689540306 |
| 7 | 0.873080032 | **1.053047968** |
| 8 | 0.247372676 | 0.930129913 |
| 9 | 0.414713015 | **1.55221519** |

Table S14:

| | TCR | HLA |
|---|---|---|
| 1 | **2.052117264** | **1.651090343** |
| 2 | **2.540716612** | **1.500519211** |
| 3 | 0.749185668 | 0.970924195 |
| 4 | **1.335504886** | 0.425752856 |
| 5 | 0.195439739 | 0.57113188 |
| 6 | **2.052117264** | 0.420560748 |
| 7 | 0.03257329 | 0.763239875 |
| 8 | 0.912052117 | **1.033229491** |
| 9 | 0.06514658 | **1.033229491** |
| 10 | 0.06514658 | **1.630321911** |

Table S15:

| | TCR | HLA |
|---|---|---|
| 1 | **2.588235294** | **1.76171875** |
| 2 | 0 | **1.375** |
| 3 | 0 | **1.24609375** |
| 4 | **1.294117647** | 0.47265625 |
| 5 | **4.529411765** | 0.544270833 |
| 6 | 0.647058824 | 0.272135417 |
| 7 | 0.647058824 | 0.272135417 |
| 8 | 0.647058824 | 0.458333333 |
| 9 | 0 | **1.002604167** |
| 10 | 0.647058824 | **1.217447917** |
| 11 | 0 | **2.377604167** |

Table S16:

| | TCR | HLA |
|---|---|---|
| 1 | 0 | **1.028571429** |
| 2 | 0 | **1.885714286** |
| 3 | 0 | **1.542857143** |
| 4 | 0 | **1.2** |
| 5 | 0 | 0.171428571 |
| 6 | **1.2** | **1.2** |
| 7 | **2.4** | 0.171428571 |
| 8 | 6 | 0 |
| 9 | 0 | 0.514285714 |
| 10 | **2.4** | **1.028571429** |
| 11 | 0 | **1.028571429** |
| 12 | 0 | **2.228571429** |

Table S17:

| | TCR | HLA |
|---|---|---|
| 1 | 0 | **1.702875399** |
| 2 | 0 | **1.495207668** |
| 3 | 0 | **1.827476038** |
| 4 | 0 | 0.747603834 |
| 5 | 0 | 0.706070288 |
| 6 | **13** | 0.581469649 |
| 7 | 0 | 0.249201278 |
| 8 | 0 | 0.041533546 |
| 9 | 0 | 0.083067093 |
| 10 | 0 | 0.664536741 |
| 11 | 0 | **1.038338658** |
| 12 | 0 | **1.24600639** |
| 13 | 0 | **2.616613419** |

#### 4.2 Interpretability based on HLA amino acid sites of different alleles

Please refer to **SI_hla_heatmap.csv** for detailed information.
